# Positive selection analyses identify a single WWE domain residue that shapes ZAP into a super restriction factor

**DOI:** 10.1101/2023.11.20.567784

**Authors:** Serina Huang, Juliana Girdner, LeAnn P Nguyen, David Enard, Melody MH Li

**Affiliations:** Department of Human Genetics, David Geffen School of Medicine, University of California, Los Angeles, CA, USA; Department of Chemistry and Biochemistry, University of California, Los Angeles, CA, USA; Department of Microbiology, Immunology and Molecular Genetics, University of California, Los Angeles, CA, USA; Molecular Biology Institute, University of California, Los Angeles, Los Angeles, CA, USA; Department of Ecology and Evolutionary Biology, University of Arizona, Tucson, AZ, USA; AIDS Institute, David Geffen School of Medicine, University of California, Los Angeles, CA, USA

## Abstract

The host interferon pathway upregulates intrinsic restriction factors in response to viral infection. Many of them block a diverse range of viruses, suggesting that their antiviral functions might have been shaped by multiple viral families during evolution. Virus-host conflicts have led to the rapid adaptation of viral and host proteins at their interaction hotspots. Hence, we can use evolutionary genetic analyses to elucidate antiviral mechanisms and domain functions of restriction factors. Zinc finger antiviral protein (ZAP) is a restriction factor against RNA viruses such as alphaviruses, in addition to other RNA, retro-, and DNA viruses, yet its precise antiviral mechanism is not fully characterized. Previously, an analysis of 13 primate ZAP identified 3 positively selected residues in the poly(ADP-ribose) polymerase-like domain. However, selective pressure from ancient alphaviruses and others likely drove ZAP adaptation in a wider representation of mammals. We performed positive selection analyses in 261 mammalian ZAP using more robust methods with complementary strengths and identified 7 positively selected sites in all domains of the protein. We generated ZAP inducible cell lines in which the positively selected residues of ZAP are mutated and tested their effects on alphavirus replication and known ZAP activities. Interestingly, the mutant in the second WWE domain of ZAP (N658A) is dramatically better than wild-type ZAP at blocking replication of Sindbis virus and other ZAP-sensitive alphaviruses due to enhanced viral translation inhibition. The N658A mutant inhabits the space surrounding the previously reported poly(ADP-ribose) (PAR) binding pocket, but surprisingly has reduced binding to PAR. In summary, the second WWE domain is critical for engineering a super restrictor ZAP and fluctuations in PAR binding modulate ZAP antiviral activity. Our study has the potential to unravel the role of ADP-ribosylation in the host innate immune defense and viral evolutionary strategies that antagonize this post-translational modification.

**Author summary:** Host proteins and viral proteins that encounter one another are locked in a perpetual genetic arms race. In this evolutionary race, a mutation that confers a survival advantage will become more frequent in the population. By looking at the sequences of genes that are known to have antiviral roles in mammals, we can identify the exact sites where a host and viral protein have interacted and gain insight into how an antiviral protein works. Here, we identified these sites in zinc finger antiviral protein (ZAP), a host protein that blocks many different viruses. We found that changing one of the sites from the original amino acid to another dramatically improves ZAP’s antiviral activity against Sindbis virus, an alphavirus, due to improved inhibition of viral translation. Our mutation is also better at inhibiting other members in the *Alphavirus* genus. We observed that our mutant ZAP has reduced ability to bind poly(ADP-ribose), a post-translational modification that is targeted by alphaviruses for productive infection. Our findings help us better understand how viruses have shaped the evolution of broad-spectrum host antiviral proteins, with great implications for the engineering of super restriction factors.

## Introduction

Viral and host proteins are constantly engaging in genetic conflicts that create selective pressures on the other side to evolve. In a host innate immune protein, an advantageous mutation that successfully maintains recognition of a viral protein or evades a viral antagonist will rise in frequency, a phenomenon called positive selection. The amino acid sites on which positive selection have acted can be identified by bioinformatic approaches when the non-synonymous substitution rate is estimated to exceed the synonymous substitution rate (1,2). The signatures of positive selection on a protein can inform us about historical interaction hotspots between the host and virus (3), as well as highlight sites that have important antiviral roles in winning the host-virus arms race.

Signatures of positive selection are especially prevalent in host interferon (IFN)-stimulated genes (ISGs) that are induced to counteract viral infections (3). One of these ISGs is zinc finger antiviral protein (ZAP), also known as poly(ADP-ribose) polymerase 13 (PARP13) (4). ZAP inhibits a diverse range of virus genera, yet its antiviral activity can be specific to particular members in a genus, suggesting viral evasion or antagonism of ZAP inhibition (5,6). For example, ZAP blocks many species of mosquito-borne alphaviruses to varying degrees, where Sindbis virus (SINV) and Ross River virus (RRV) are more sensitive than o’nyong’nyong virus (ONNV) and chikungunya virus (CHIKV) vaccine strain 181/clone 25 (7,8). Alphaviruses have a positive-sense RNA genome, which can be immediately translated into viral proteins by host ribosomes upon entry into the host cell (9,10). The viral proteins then replicate the viral genome, leading to the production of structural proteins and the assembly of mature virus particles. It is in the early stages of infection that ZAP acts to prevent the translation of alphaviral RNA by synergizing with the host E3 ubiquitin ligase, tripartite motif containing 25 (TRIM25) (11,12).

ZAP has two major splice isoforms, ZAPS (short) and ZAPL (long), with distinct antiviral and immunomodulatory activities (7,13–15). Recently discovered isoforms, ZAPM (medium) and ZAPXL (extralong), resemble the antiviral activities of ZAPS and ZAPL, respectively (7). The N-terminus of ZAP contains four zinc fingers (ZnFs) that bind RNA. It is followed by a fifth ZnF and two WWE domains, named for its motif containing tryptophan, tryptophan, and glutamic acid. The ADP-ribose-binding ability of the second WWE domain (WWE2) has only been recently discovered (16,17). At the C-terminus, there is a PARP-like domain. Despite being one of the 17 PARPs, ZAP is the only PARP with a PARP-like domain that is catalytically inactive and cannot ADP-ribosylate substrates (18,19), but confers more antiviral activity on the longer isoforms (7,15,20,21). Even though the RNA binding activity of ZAP has been extensively studied, how the other domains contribute to ZAP antiviral activity are not well characterized.

While ZAP has been shown to be positively selected (15,22), there are outstanding questions about the antiviral mechanism of ZAP and how its cellular functions contribute to viral inhibition. A previous study performed positive selection analysis on ZAP sequences from 13 primate species and found 3 positively selected sites, all in the PARP-like domain. We aimed to expand upon this study because ZAP is effective against diverse groups of viruses including but not limited to lentiviruses. ZAP has a long-standing history of host-virus interactions and likely arose from a gene duplication event after the divergence of tetrapods (23). Assuming that at least some of the positively selected sites are driven by the ancestors of extant ZAP-sensitive viruses (e.g. alphaviruses, flaviviruses, coronaviruses, etc.), we would expect to detect positive selection signals from a broader range of mammals which these viruses tend to infect.

Here, we performed positive selection analyses on 261 mammalian ZAP sequences using four complementary and sophisticated models that make more realistic assumptions about the substitution rates. We identified 7 residues that are positively selected in ZAP, most of which are outside the PARP-like domain. We mutated each positively selected site and found that one mutant in the WWE2 (N658A) has antiviral activity that is almost 10 times stronger than wild-type (WT) ZAP against SINV, creating a super restrictor that is more antiviral than any versions of ZAP that were previously reported. The N658A mutant is more efficient than ZAPL WT at inhibiting virion production of SINV and replication of a panel of alphaviruses in a manner that is dependent on viral translation suppression. Interestingly, mutation of both positively selected sites in the WWE2 that form a potential interaction surface does not further increase the antiviral activity of ZAP.

We then investigated the role of viral RNA binding, TRIM25 interaction, IFN response, and poly(ADP-ribose) (PAR) binding in mediating the activity of a super restrictor ZAP. We found that the superior antiviral activity of the N658A mutant can be attributed to changes in PAR binding by the ZAPL mutant. We mutated site 658 to orthologous residues found in other mammalian species and observed that all of them improved ZAP inhibition against SINV. This surprising finding suggests that evolutionary forces did not steer human ZAP to be the most antiviral, at least not against alphaviruses. By taking into account the history of host-virus conflicts, positive selection analyses allow us to identify specific sites with high impact on the effectiveness of the host antiviral program, providing a blueprint for generating super restriction factors.

## Results

### ZAP is positively selected throughout mammalian evolution at novel sites

We used the longest isoform of ZAP, ZAPXL, to curate and align 261 high quality mammalian orthologs. We ran 4 positive selection tests with complementary strengths on the alignment of mammalian ZAP sequences: Fixed Effects Likelihood (FEL); Mixed Effects Model of Evolution (MEME); Fast, Unconstrained Bayesian AppRoximation (FUBAR); and the Bayesian mutation-selection model by Rodrigue *et al* (24–27). FEL does not make assumptions about the distribution of selection parameters over sites but assigns independent nonsynonymous and synonymous rates to each site. MEME accounts for the fact that positive selection occurs episodically, rather than remaining constant over time. FUBAR improves upon random effect likelihood models (28) by implementing more parametrically complex models. Rodrigue *et al.*’s method is the first Bayesian mutation-selection model, offering higher sensitivity.

To validate the robustness of our tests, we ran the 13 primate ZAP sequences from the study by Kerns *et al.* and were able to replicate the 3 positively selected sites previously identified. Using the 261 mammalian ZAP, we identified 7 positively selected sites that are shared by all 4 tests (S1A Fig) and mapped them to human ZAP isoforms (S1B Fig). For consistency, the positively selected sites are numbered in the context of ZAPL and ZAPS, which are the more well studied isoforms with antiviral activities similar to ZAPXL and ZAPM, respectively. The positively selected sites we identified are concentrated in specific regions spanning across the ZAP gene (Fig 1A). Two of these sites are within the first 254 amino acids of the protein, which comprise the RNA binding domain that is necessary for ZAP recognition and inhibition of viral RNA. These residues, Q28 and C38, are relatively close to each other but are positioned opposite the RNA binding groove, with both of their side chains pointing away from the rest of the structure (29) (Fig 1B). We had not expected any sites to be in the RNA binding domain because RNA binding is an essential function of ZAP. However, the identification of these two sites raises the possibility that viral proteins can interact with ZAP at a different location in its N-terminal region without interfering with binding to viral RNA.

**Figure 1.**
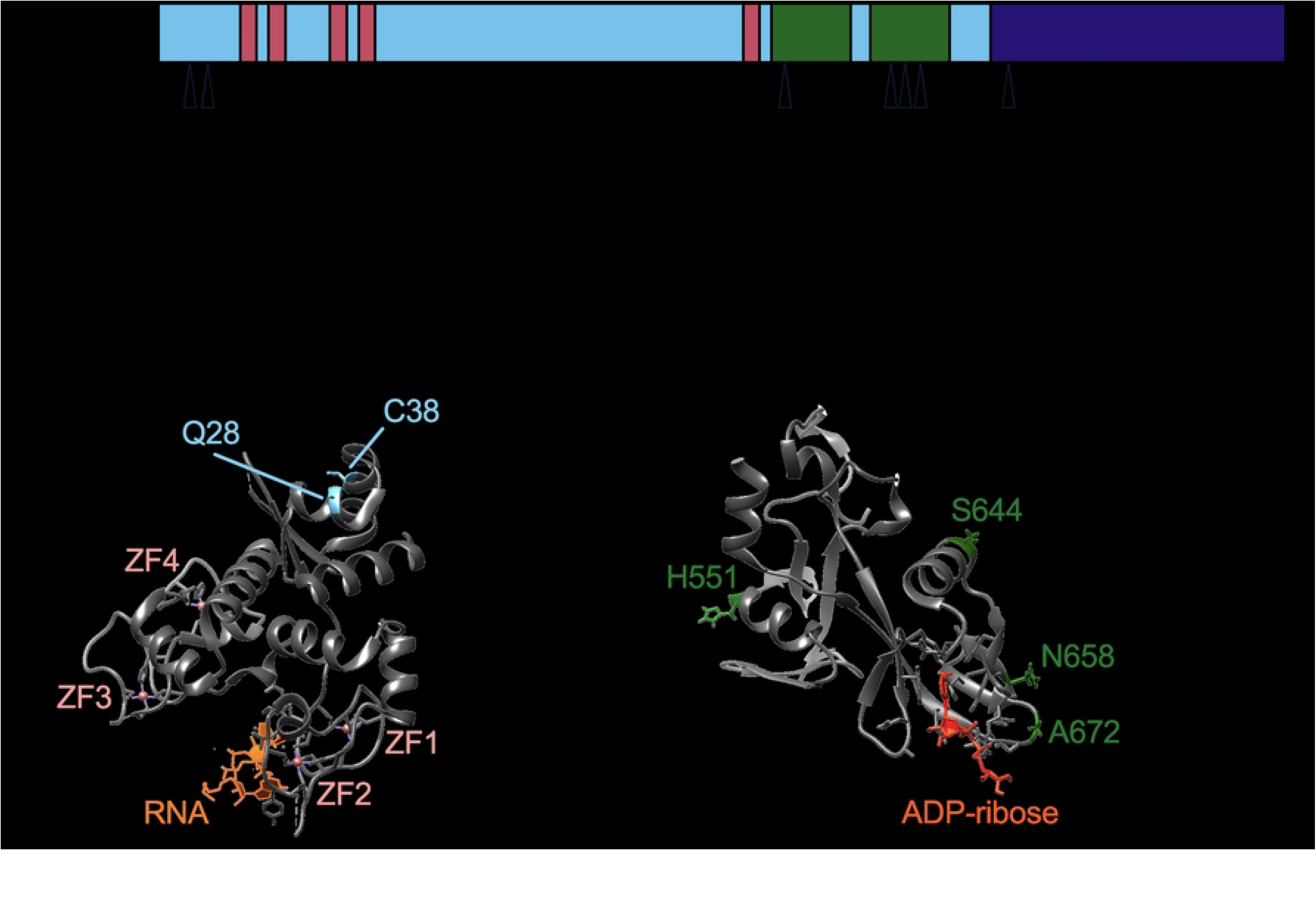
Identification of 7 positively selected sites across ZAP protein. (A) A schematic of the ZAPL isoform annotated with its domains. Triangles indicate positively selected sites identified from the overlap of four methods: Fixed Effects Likelihood; Mixed Effects Model of Evolution; Fast, Unconstrained Bayesian AppRoximation; and the Bayesian mutation-selection model by Rodrigue *et al*. (B) ZAP RNA binding domain bound to RNA. The structure (PDB: 6UEJ) (29) is visualized with UCSF Chimera. Positively selected Q28 and C38 residues shown in blue; RNA in orange; zinc fingers in salmon. (C) ZAP central domain bound to ADP-ribose. The structure (PDB: 7TGQ) (17) is visualized with UCSF Chimera. Positively selected sites H551, S644, N658, and A672 shown in green; ADP-ribose in dark orange.

More than half of the positively selected sites are in the central domain, 3 of which are tightly clustered in the WWE2, which has only recently been found in ZAP to bind PAR. When mapped to the available crystal structure of the central region consisting of the fifth zinc finger and the two WWE domains (16,17), two of the sites, N658 and A672, are next to the PAR binding pocket and face outward, supporting that they are at the interface of host-virus interactions (Fig 1C). Taken together, our positive selection analyses demonstrate that ZAP has been rapidly evolving not just during primate evolution, but also during mammalian evolution. These novel positively selected residues in ZAP are found in all domains of ZAP, suggesting that ancient viruses have likely targeted and antagonized ZAP at distinct sites.

### Most of the positively selected sites affect ZAP antiviral phenotype against SINV

To probe the effect of the positively selected sites, we mutated each site from the WT amino acid in humans to alanine because alanine is chemically inert and would not dramatically change the secondary structure of the protein (30). In the case where the WT amino acid is alanine, we mutated it to valine, the next closest amino acid. We cloned either WT or mutant ZAPS and ZAPL with an N-terminal FLAG tag into the ePiggyBac (ePB) transposon system and generated stable cell lines in ZAP knockout (KO) HEK293T cells (31,32). We tested the mutants in the ZAPS and ZAPL background because ZAPS and ZAPL are most commonly studied and have comparable antiviral activities to ZAPM and ZAPXL, respectively.

Almost all the mutant cell lines have robust ZAP expression when induced by doxycycline, with the exception of ZAPS Q28A which appears to have a truncation at the C-terminus, as it is still able to be detected by the N-terminal FLAG tag (Fig 2A). Since our candidate sites are positively selected throughout mammalian evolution, we chose to test their antiviral activity against alphaviruses, whose primary hosts are mammals such as primates, horses, and rodents (33). We first infected the ZAP cell lines with SINV, a prototype alphavirus that is susceptible to ZAP inhibition.

**Figure 2.**
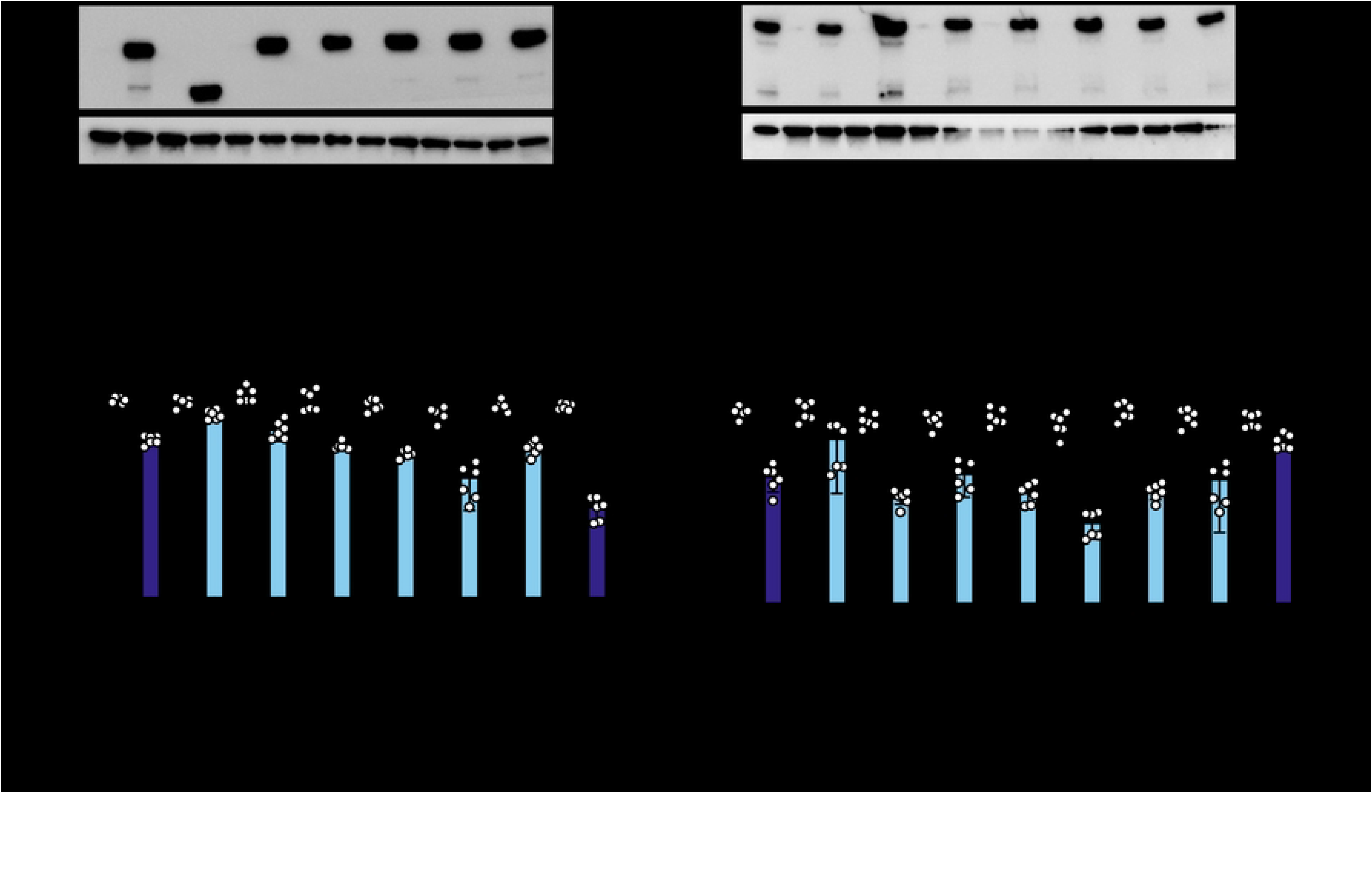
ZAP mutated at positively selected sites show differential antiviral activity against SINV. (A, C) Western blot of (A) ZAPS or (C) ZAPL wild-type (WT) or positive selection mutants inducible ZAP KO 293T cell lysates. (B, D) (B) ZAPS or (D) ZAPL WT or mutant ZAP KO 293T cells were induced for ZAP 24 hours before infection with SINV Toto1101/Luc at a multiplicity of infection (MOI) of 0.01 plaque forming units (PFU)/cell and harvested at 24 hours post-infection (h.p.i.) for luciferase assay by measuring relative luciferase units (RLU). Data are combined from two independent experiments; error bars indicate standard deviation. 1µg/mL dox is used to induce ZAP expression. Asterisks indicate statistically significant differences as compared to the -dox condition for each cell line (Two-way ANOVA and Tukey’s multiple comparisons test: *, p<0.05; **, p<0.01; ***, p<0.001; ****, p<0.0001).

We infected ZAPS and ZAPL WT and mutant cell lines with a luciferase-expressing SINV reporter virus. Despite differences in absolute fold inhibitions between independent experiments featuring ZAPS and ZAPL mutants, we found that ZAPL WT is invariably more antiviral than ZAPS WT, consistent with previous reports (7,15). While a couple of mutants are considerably less antiviral than the corresponding WT ZAP (ZAPS Q28A and C38A; ZAPL Q28A and H551A), most mutants have enhanced antiviral activity, as evidenced by a greater fold inhibition than the corresponding WT ZAP (ZAPS mutants ranging from 16- to 123-fold vs. 12-fold for ZAPS WT; and ZAPL mutants ranging from 103- to 580-fold vs. 78-fold for ZAPL WT). These results suggest that most PS sites are not evolutionarily optimized for anti-SINV activity in humans and *could* be enhanced (Fig 2B & Fig 2D). Notably, the N658A mutant located in the WWE2 shows a significant improvement in ZAP antiviral activity (10 times better than ZAPS WT and 7 times better than ZAPL WT). In addition, some mutants displayed isoform-specific effects. For instance, ZAPL C38A is more antiviral than ZAPL WT, but its ZAPS counterpart is less antiviral than ZAPS WT. These results suggest that altering the WT amino acid at these positively selected sites *a posteriori* changes the antiviral activity of ZAP against SINV—many stronger than before—and that adaptations at these sites have important functional consequences.

Since both sites 658 and 672 are located in the WWE2 and flank the PAR binding pocket in the crystal structure (Fig 1A & 1C), we wondered if the two sites constitute a critical host-virus interface in concert, as is the case with TRIM5α (34). We generated the double mutant N658A/A672V (NA) in the same ZAP KO ePB system and assessed its ability to restrict SINV replication. Both ZAPS and ZAPL NA double mutants are as stably expressed as the single mutants (Fig 3A & 3C). To our surprise, the antiviral activity of the ZAPS NA double mutant is not an intermediate between ZAPS N658A (33x) and A672V (5x); rather, it reduces the antiviral activity of N658A to that of ZAPS WT and A672V (Fig 3B, 4 to 5x), suggesting that A672V has a dominant negative effect on N658A in ZAPS. The ZAPL NA double mutant increases antiviral activity against SINV replication by less than 2-fold (103x vs. 58x for ZAPL WT). However, it does not approach the strength of ZAPL N658A (224x) and A672V (225x) (Fig 3D). The differential antiviral activity of the A672V single mutant and the NA double mutant in ZAPS and ZAPL again highlights isoform specificity at particular sites. Together, the WWE2 mutations in combination lessen the increase in antiviral activity we observed with the single N658A mutation in both ZAPS and ZAPL backgrounds, suggesting that these mutations may not act as a single protein interaction surface.

**Figure 3.**
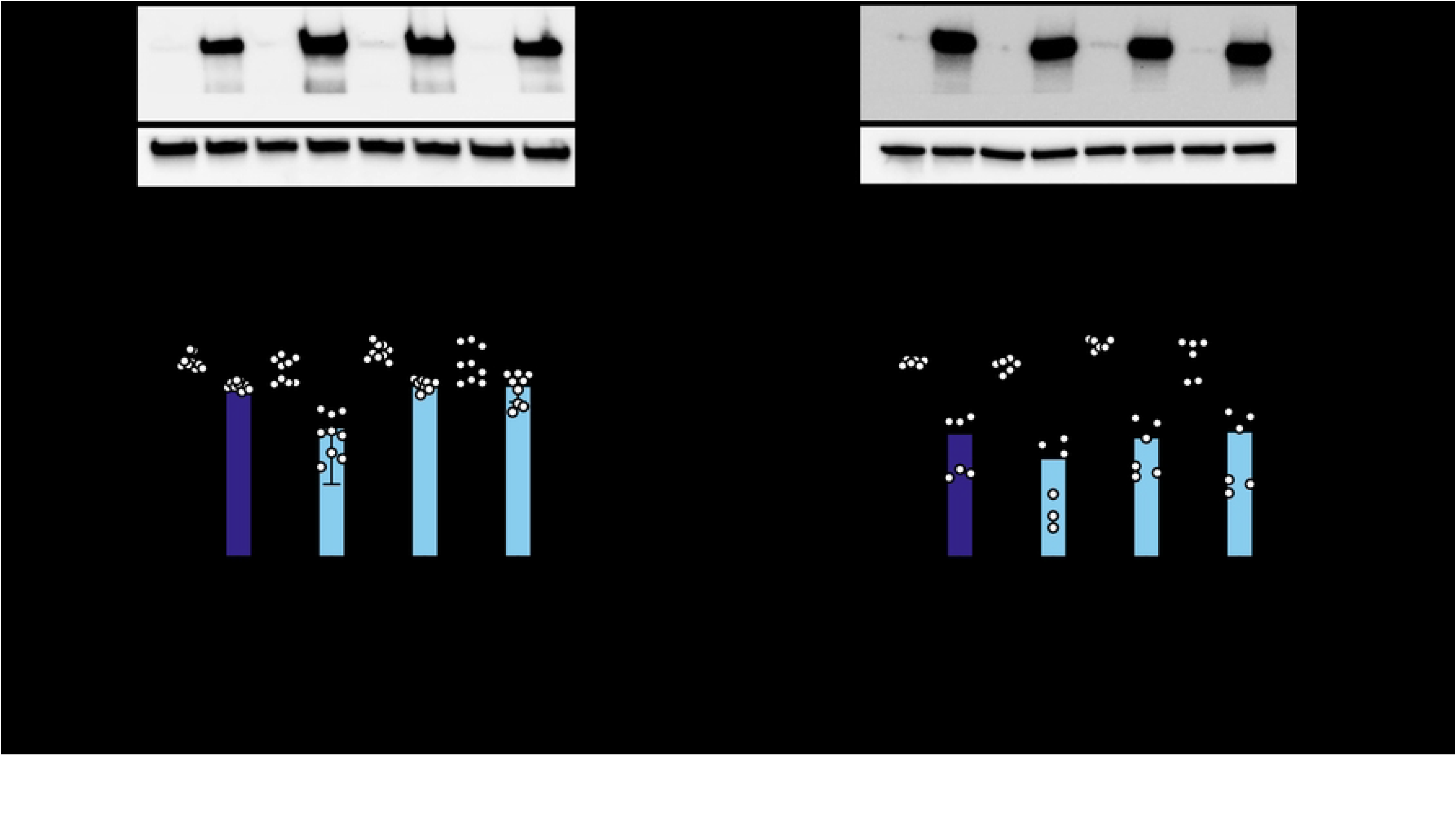
Mutating both positively selected sites in the second WWE domain of ZAP does not further enhance antiviral activity. (A, C) Western blot of (A) ZAPS or (C) ZAPL WT, N658A, A672V, or N658A/A672V (NA) double mutant inducible ZAP KO 293T cell lysates. (B, D) (B) ZAPS or (D) ZAPL WT or mutant ZAP KO 293T cells were induced for ZAP expression 24 hours before infection with SINV Toto1101/Luc at an MOI of 0.01 PFU/cell and harvested at 24 h.p.i for luciferase assay. Data are combined from three (B) and two (D) independent experiments; error bars indicate standard deviation. 1µg/mL dox is used to induce ZAP expression. Asterisks indicate statistically significant differences as compared to the -dox condition for each cell line (Two-way ANOVA and Tukey’s multiple comparisons test: **, p<0.01; ****, p<0.0001).

### The ZAPL N658A mutant blocks the early steps of alphaviral infection more effectively

We were interested by the superior antiviral activity of the N658A mutant alone and focused on the ZAPL isoform to study the mutant in the presence of all domains of ZAP, including the PARP-like domain. We wanted to determine whether the effects on viral replication impact the overall virion production. We infected ZAPL WT or N658A cells with SINV and collected the cell supernatant containing mature and released virions at 0, 6, 12, 24, and 36 hours post-infection (h.p.i.). We determined the viral titer on BHK-21 cells via plaque assay. We found that both ZAPL WT and N658A significantly inhibited SINV virion production, but at 24 h.p.i., ZAPL N658A is about 4-fold more inhibitory (Fig 4A, 11x vs. 40x), consistent with the phenotype we observed with viral replication.

**Figure 4.**
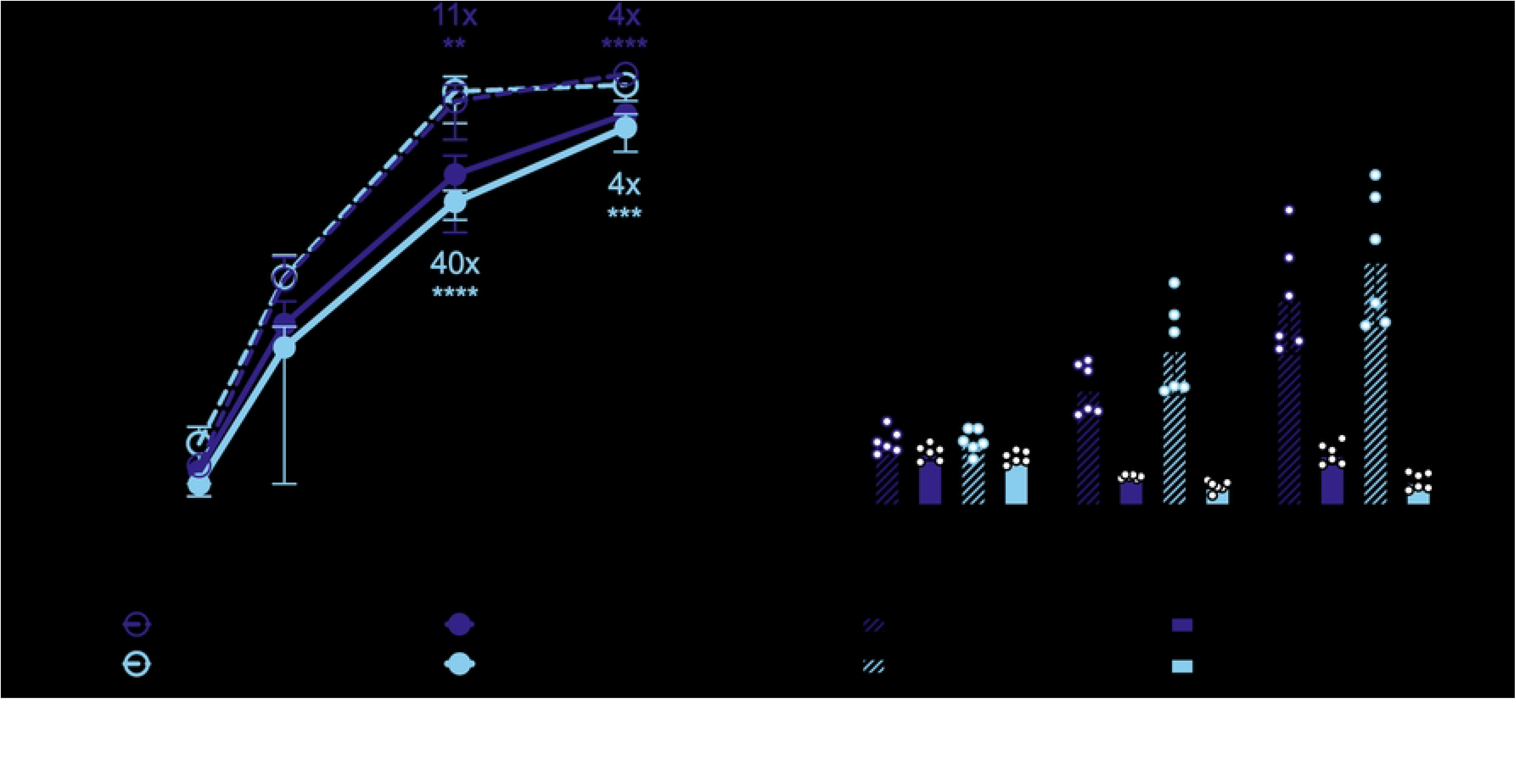
The N658A mutant is better at inhibiting virion production and SINV RNA translation. ZAPL WT or N658A ZAP KO 293T cells were induced for ZAP expression with 1μg/mL dox 24 hours prior to infection. Cells were infected with (A) SINV Toto1101 at an MOI of 0.01 PFU/cell, harvesting supernatant at 6, 12, 24, 36, and 40 h.p.i. for plaque assays. Viral titers of plaque assays are determined in BHK-21 cells. Data are combined from two independent experiments; error bars indicate standard deviation. Asterisks indicate statistically significant differences as compared to the -dox condition; or (B) SINV Toto1101/Luc:ts6 at an MOI of 0.01 PFU/cell, and harvested at 0, 3, and 6 h.p.i. for luciferase assay. Data are combined from two independent experiments; error bars indicate standard deviation. Asterisks indicate statistically significant differences as compared to the -dox condition for each cell line (Fisher’s Least Significant Difference Test (A) or Two-way ANOVA and Tukey’s multiple comparisons test (B): **, p<0.01; ***, p<0.001; ****, p<0.0001).

Next, we sought to determine the stage in the viral life cycle at which the ZAPL N658A mutant acts. Because ZAP is known to act by blocking alphaviral RNA translation, we tested the positively selected ZAP mutant N658A against a temperature-sensitive replication-deficient SINV luciferase reporter virus that cannot replicate at 40°C (35). We infected ZAP WT and N658A cell lines with the replication-deficient virus at the non-permissive temperature and found that the N658A mutant is better at blocking SINV RNA translation (Fig 4B). Our finding supports that the superior antiviral activity of the N658A mutant is likely due to an enhanced block at the step of incoming viral RNA translation.

Since we hypothesized that the positive selection of ZAP may be driven by ancient alphavirus-like viruses, we tested whether the N658A mutant also inhibits other alphaviruses better. We infected the ZAPL WT or N658A cell line with GFP-expressing SINV, RRV, ONNV, CHIKV vaccine strain 181/clone 25, and VEEV. Alphaviruses known to be more sensitive to ZAP inhibition are more inhibited by the N658A mutant (Fig 5A, 5x vs. 38x against SINV; Fig 5B, 24x vs. 132x against RRV), while the ones that are less sensitive were similarly resistant to both ZAPL WT and N658A (Fig 5C, 3.8x vs. 8.3x against ONNV; Fig 5E, 1.1x vs. 1.1x against VEEV). Interestingly, even though we previously observed that the non-reporter CHIKV vaccine strain is less susceptible to ZAP inhibition (7), we saw that both ZAPL WT and N658A dramatically inhibited GFP-expressing CHIKV vaccine strain, with the N658A mutant being more antiviral than WT (Fig 5D). Since the CHIKV strain we tested expresses the GFP reporter under the control of the viral subgenomic promoter, our results suggest that ZAP might inhibit step(s) at or prior to viral subgenomic mRNA expression.

**Figure 5.**
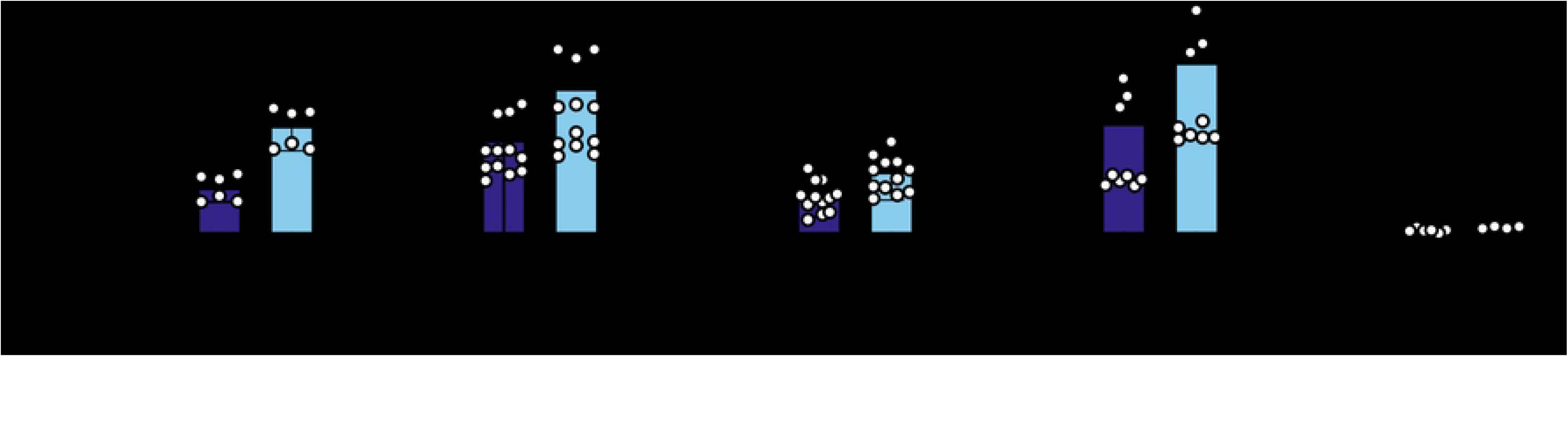
The ZAPL N658A mutant inhibits many other alphaviruses better than WT. After 24 hours of 1µg/mL dox treatment, ZAPL WT or N658A ZAP KO 293T cells were infected with (A) GFP-expressing Sindbis virus (SINV, MOI = 0.01), (B) Ross River virus (RRV, MOI = 1), (C) o’nyong’nyong virus (ONNV, MOI = 1), (D) chikungunya virus (CHIKV, MOI = 0.1), or (E) Venezuelan equine encephalitis virus (VEEV, MOI = 0.1) PFU/cell for 24 hours before their percentage of infection was determined by flow cytometry. Data are combined from at least two independent experiments of biological replicates in triplicate wells; error bars indicate standard deviation.

### The improved antiviral activity of the N658A mutant is not due to changes in binding to SINV RNA, interaction with TRIM25, or increased activation of ISGs

To determine the mechanism of the enhanced antiviral activity of the N658A mutant, we characterized the mutant in terms of known abilities of ZAP. Since ZAP is recognized as a sensor of CpG-rich viral RNA, we wondered if N658A binds better to SINV genomic RNA than ZAPL WT does. We performed an *in vitro* RNA pulldown assay by incubating protein lysates from either the ZAPL WT or N658A cell line with equal amounts of biotinylated SINV genomic RNA. We pulled down the biotinylated viral RNA using streptavidin beads and probed for ZAP. We generated and tested a ZAP KO HEK293T cell line with inducible expression of a ZAPS C86A/Y96A mutant (ZAPS CY), which is deficient in RNA binding (36,37), as negative control. As expected, markedly less ZAPS CY is bound to equal amounts of SINV RNA compared to ZAPL WT. Equal amounts of ZAPL WT and ZAPL N658A are bound to SINV RNA (Fig 6A), suggesting that factors other than viral RNA binding may contribute to the enhanced antiviral activity of the mutant.

**Figure 6.**
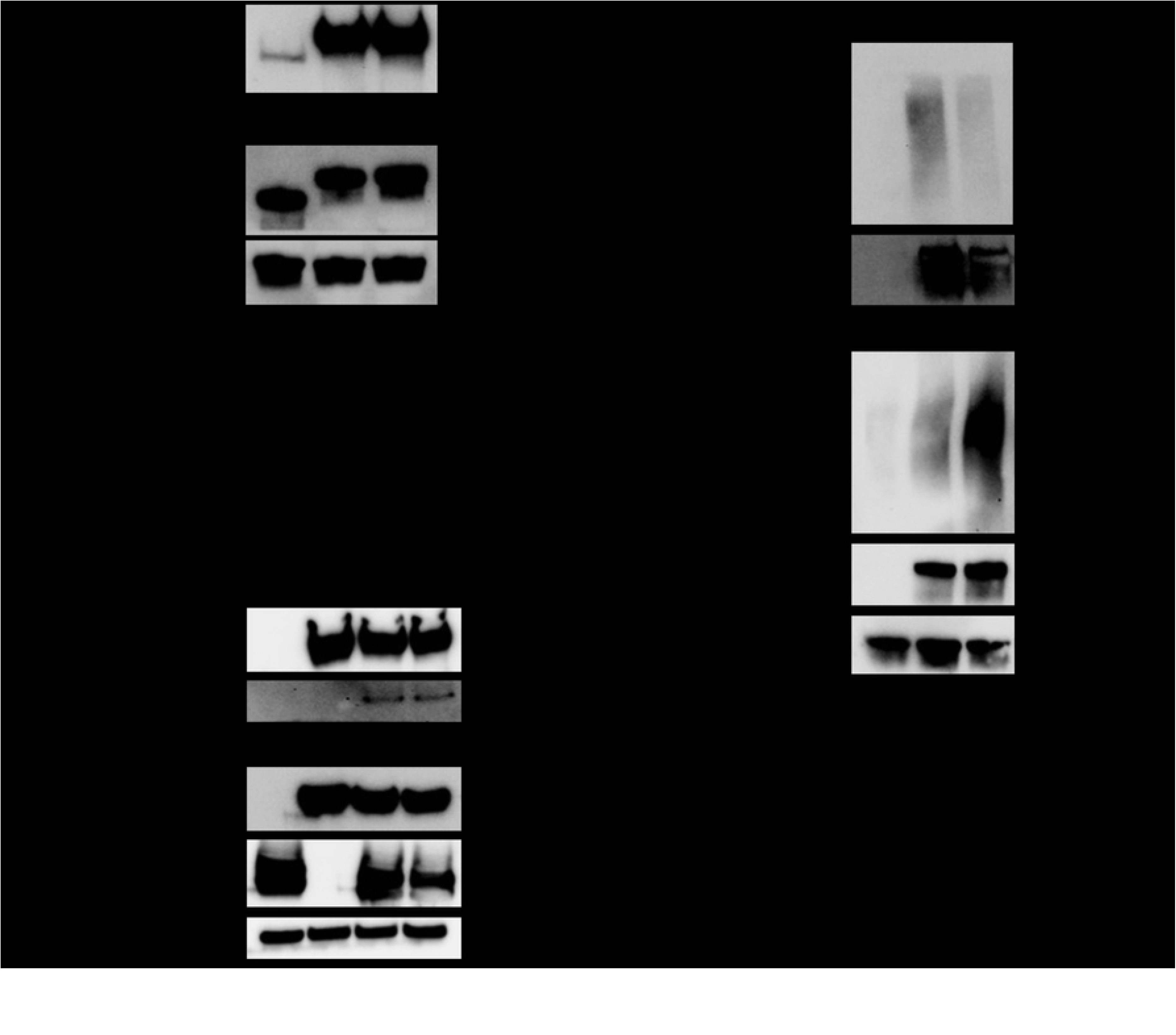
The improved antiviral activity of the N658A mutant is not due to changes in binding to SINV RNA or interaction with TRIM25, but changes in binding to poly(ADP-ribose) (PAR) (A) Western blot of ZAPL CY, WT, or N658A inducible ZAP KO 293T cell lysates bound to biotinylated SINV genomic RNA immunoprecipitated by streptavidin beads. Data are representative of three independent experiments. (B) Western blot of ZAP KO 293T transfected with pcDNA3.1-3XFLAG-ZAPL and pcDNA3.1-myc-TRIM25. Lysates were immunoprecipitated by FLAG beads. Data are representative of four independent experiments. (C) Western blot of ZAPL WT or N658A inducible ZAP KO 293T cell lysates immunoprecipitated by FLAG beads after treatment with 1µM PARG inhibitor. Data are representative of three independent experiments. 1µg/mL dox is used to induce ZAP expression in ePB ZAP inducible cell lines.

We then asked whether the N658A mutant changes ZAP’s ability to interact with TRIM25, a host E3 ubiquitin ligase that is a requisite cofactor for ZAP’s inhibition of viral RNA translation (11,12). We transfected 3XFLAG-ZAPL and myc-TRIM25 into ZAP KO HEK293T cells and performed a co-immunoprecipitation with FLAG beads. We found that ZAPL WT and N658A interact with TRIM25 similarly (Fig 6B), suggesting that the increased antiviral activity of the N658A mutant is not related to changes to its synergy with TRIM25.

We further evaluated whether increased IFN induction is responsible for the enhanced antiviral activity of the ZAPL N658A mutant. After treating ZAPL WT and N658A cell lines with poly(I:C), a double-stranded RNA mimic, to stimulate the IFN response, we performed quantitative PCR analysis of the mRNA levels of IFN-β and IFIT1, a classical antiviral ISG. We found that poly(I:C) treatment upregulates IFN-β and IFIT1 levels, and expression of ZAPL WT and N658A further augments the response (S2 Fig). Importantly, both IFN-β and IFIT1 induction levels in ZAPL WT and N658A cell lines are similar upon stimulation (S2 Fig), ruling out a heightened IFN response as responsible for the N658A improved antiviral phenotype.

### The ZAPL N658A mutant has reduced binding to PAR

Since RNA binding, TRIM25 interaction, and the IFN response do not appear to mediate the superior antiviral activity of ZAPL N658A, we decided to characterize the effects of the mutation on WWE domain function. The WWE2 in ZAP has recently been found to bind to PAR, an ability that enhances ZAP’s antiviral function against a CpG-enriched HIV-1 (17). We wondered if mutating site 658, which is within the WWE2, changes ZAP’s ability to bind to PAR. We performed a co-immunoprecipitation assay in which we pulled down ZAP and probed for PAR. PAR levels in the whole cell lysate are markedly lower in cells without ZAP induced (Fig 6C). Compared to ZAPL WT, ZAPL N658A binds to less PAR (Fig 6C). Altogether, these data suggest that an alanine mutation at site 658 negatively impacts ZAPL’s ability to bind PAR, despite the site being outside of the PAR binding groove. The mutation might prevent an active PARP from accessing and PARylating ZAPL in an uninfected cell. Contrary to the Q668R mutation in the PAR binding pocket which diminishes ZAP PAR binding and anti-HIV activity (17), our N658A mutant is less proficient in binding PAR, but surprisingly more adept at restricting SINV.

### Asparagine is the predominant amino acid at site 658 in ZAP yet the least antiviral

To further understand the requirements at site 658 for ZAP to become a super restrictor, we analyzed the amino acid distribution in our mammalian ZAP sequences. We observed that site 28, one of the positively selected sites, displays an even distribution of amino acids (Fig 7A). In contrast, at site 658, asparagine is the most prevalent amino acid in our 261 mammalian ZAP sequences (68%, Fig 7A). In terms of the amino acid property, there is less variation at site 658 than at site 28. Even though site 658 has rapidly evolved, polar amino acids seem to be favored by evolution. 80% of the mammals in our alignment have a polar amino acid at site 658: 177 out of the 261 mammals (68%) have asparagine and 32 (12%) have serine (Fig 7A). This is in stark contrast to Q28, where every amino acid property is present: 7% have a nonpolar amino acid (alanine, glycine); 38% have a polar amino acid (glutamine, asparagine, serine); 38% have a negatively charged amino acid (aspartic acid, glutamic acid); and 23% have a positively charged amino acid (histidine, lysine, arginine) (Fig 7A & Fig 7C), demonstrating that site 28 is able to tolerate more flexibility in the chemical property of its amino acid. Sites that are not under positive selection, 657 and 659, show even less amino acid diversity (Fig 7B). Site 657 is dominated by a polar (glutamine) or positive (arginine) amino acid, and site 659 permits only nonpolar amino acids with an aromatic ring (tyrosine and phenylalanine).

**Figure 7.**
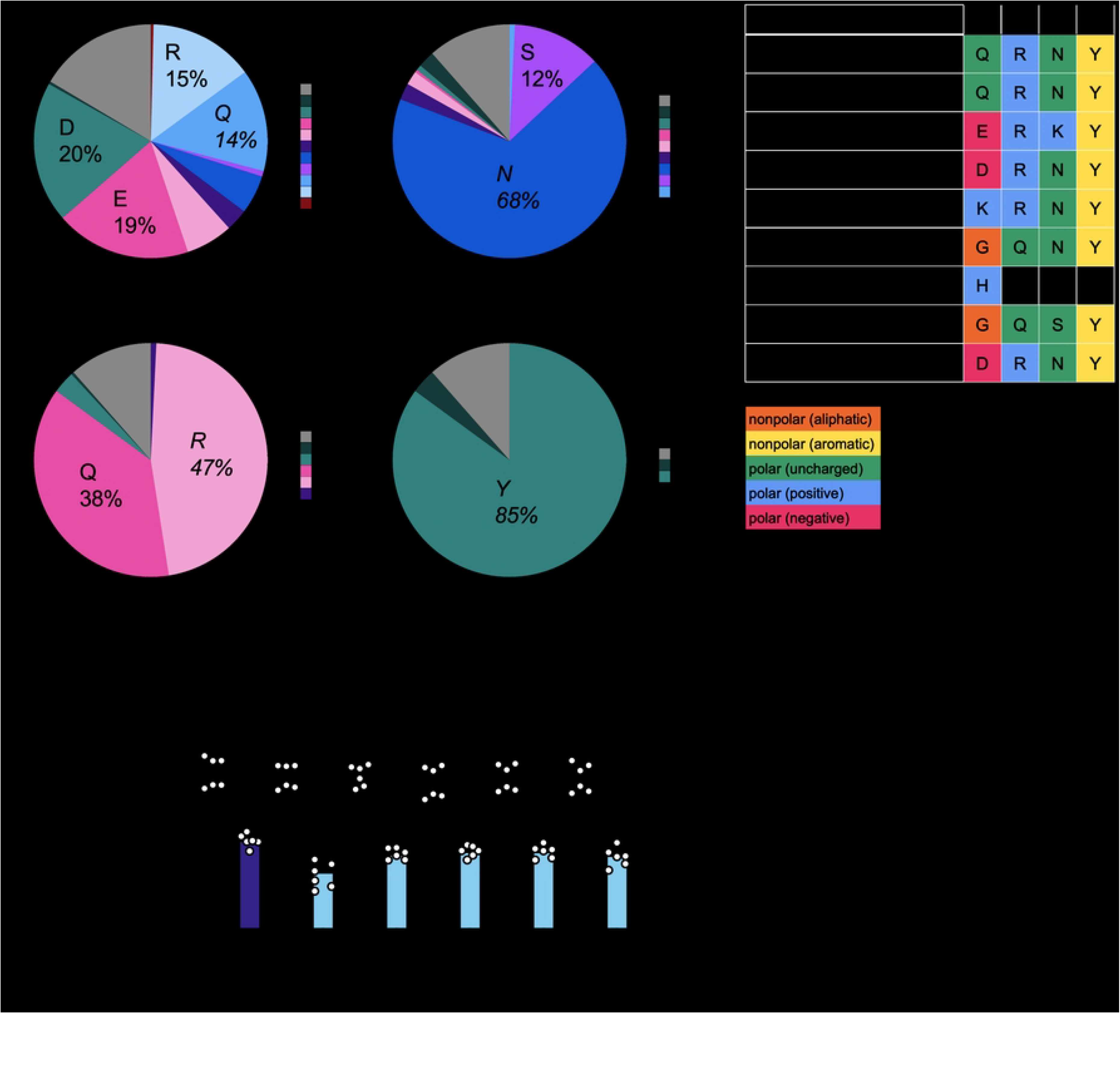
Asparagine is the predominant amino acid at site 658 yet confers weaker antiviral activity. (A, B) The distribution and (C) an abridged alignment of amino acids at sites 28, 657, 658, and 659. (D) ZAPL WT or N658A ZAP KO 293T cells were induced for ZAP expression with 1μg/mL dox. Cells were infected with SINV Toto1101/Luc at an MOI of 0.01 PFU/cell and harvested at 24 h.p.i for luciferase assay. Data are combined from two independent experiments; error bars indicate standard deviation. Asterisks indicate statistically significant differences as compared to the -dox condition for each cell line (Two-way ANOVA and Tukey’s multiple comparisons test: *, p<0.05; ****, p<0.0001).

To ascertain if a specific amino acid or a nonpolar property is required at site 658 to achieve better antiviral activity, we generated additional ZAPL N658 mutants by mutating the WT residue in humans, asparagine, to residues found in other mammalian species such as glycine (nonpolar; in African woodland thicket rat), serine (polar uncharged; in California deer mouse), lysine (positive; in greater bamboo lemur), or aspartic acid (negative; in little brown bat). We infected cell lines with inducible expression of each of these ZAPL site 658 mutants with the same luciferase-expressing SINV and found that all of them exhibit higher antiviral activity (Fig 7C), including the polar N658S, although none attains the strength of alanine mutation. Taken together, these findings suggest that just mutating asparagine to another amino acid is sufficient to improve ZAP, but the ones with the best antiviral activities (N658A and N658D) are either nonexistent or rare in nature. Further studies are required to understand why positive selection has selected for a version of ZAP that does not maximize its antiviral activity.

## Discussion

In this study, we sought other positively selected sites beyond the 3 previously identified in the PARP-like domain of ZAP and asked whether they have also been shaped into interaction interfaces during evolution. We identified 7 positively selected sites in total throughout mammalian evolution of ZAP, with only 1 residing in the PARP-like domain, supporting the notion that ZAP has been the target in more than one host-virus arms race. Notably, 4 of these positively selected sites are concentrated in the central region. We found that mutating each of these 7 positively selected sites confers differential antiviral activities against SINV. Specifically, a mutation at the WWE2 (N658A) was almost 10 times better at inhibiting SINV and other Old World alphaviruses than WT ZAP. In line with a deep mutational scanning study of TRIM5α (38), we observed that it is possible for a positively selected site to maintain strong antiviral activity when mutated to other amino acids.

Most analyses of positive selection in innate immune factors have focused on a subset of species. For example, using 17 primate TRIM5α sequences, Sawyer *et al.* identified 5 residues under positive selection all within a 13-amino acid patch that is responsible for species specificity against lentiviruses (34). Enabled by the more comprehensive sequences and robust codon substitution models presently, we hypothesized that including more species would allow us to detect positive selection signatures in regions across the whole protein and provide a more well-rounded picture of antiviral effectors. Consistent with a study that identified distinct positively selected sites in SAMHD1 using different subsets of mammals (39), we found that positively selected sites in ZAP, while concentrated, are not just restricted to the PARP-like domain (15), but span the N-terminus, central region, and C-terminus. This reflects the highly diverse and long evolutionary history of ZAP, which arose during the emergence of tetrapods (23). Further positive selection analyses in subsets of mammals are required to confirm if each positively selected site or domain is driven by distinct viruses.

We found that mutating the N658 site in the WWE2 of ZAP results in a ZAP that has stronger anti-alphavirus function yet diminishes PAR binding ability. ADP-ribosylation may be a post-translational modification exploited by alphaviruses, as a productive alphaviral infection relies on the binding to and removal of ADP-ribose by the highly conserved alphaviral macrodomains encoded by nonstructural protein 3 (40–43). It is possible that having less PAR bound to ZAPL N658A minimizes interaction between the SINV macrodomain and ZAP, thus allowing ZAP to evade recognition by a viral antagonist. Alternatively, decreased PAR binding to the ZAPL N658A mutant may also be a way to reduce PAR-dependent ubiquitination of ZAP to prevent ZAP degradation (44). Further studies are required to understand how macrodomains, PARylation, and/or ubiquitination play a role in virus infection.

Why has evolution selected for an amino acid at site 658 that makes a less antiviral version of ZAP in humans? One possibility is that having a stronger antiviral activity incurs a fitness cost on the host cell by interfering with non-immune-related cellular functions of ZAP. In cells not infected by a virus, PAR was bound to ZAP; when cells were treated with arsenite to induce stress granule formation, the amount of PAR on ZAP increased and miRNA-mediated silencing decreased (45). While the direct mRNA targets bound by ZAP and the miRNA complex remain mostly unknown, ZAP is implicated in the regulation of host transcripts in a non-viral context. A recent RNA-seq analysis also discovered that ZAPS and ZAPL bind to host mRNAs involved in the unfolded protein response and the epithelial-mesenchymal transition (46). Indeed, many genes that show strong signatures of positive selection participate in both the proper functioning of the cell and the host-virus conflict. An example gene is the Niemann-Pick C1 protein, which is an intracellular cholesterol transporter and a viral receptor for filoviruses (47,48). It would be interesting to explore if any intrinsic cellular functions of ZAP are affected by the more antiviral N658A mutation.

ZAP is a broad-spectrum antiviral protein that is effective against members from a wide range of virus families. Therefore, it is possible that some of our positively selected sites did not have a dramatically better antiviral effect compared to WT ZAP because the selection at these other sites were driven by ancient viruses that were not alphavirus-like. We wonder how our other positive selection mutants would behave against other viruses that infect mammals as their primary reservoir hosts. For instance, alphaviruses and flaviviruses share similar transmission cycles where they circulate between wild mammals and domestic mammalian dead-end hosts. Coronaviruses also commonly exploit mammals as hosts, such as camels for MERS and bats for SARS-CoV-1. If flavivirus- or coronavirus-like viruses drove the positive selection of ZAP, we expect to see a greater impact on its antiviral activity when ZAP mutants are tested against those viruses. Alternatively, viruses that are not susceptible to the increased antiviral activity of the N658A mutant might encode viral antagonists of ZAP. Notably, we saw that there was no difference in the ability of ZAPL WT and N658A to inhibit VEEV. It is possible that VEEV encodes a viral antagonist that can still recognize ZAP despite the mutation and thus is impervious to any improvement in ZAP’s antiviral activity. To determine whether the positively selected sites form an exclusive interaction interface, future studies should test more viruses from different families.

Our study is one of the first that look at positive selection of a broad-spectrum antiviral protein in a comprehensive and diverse group of mammals. By understanding what makes a super-restrictor and the host cell constraints, we can design better antiviral therapeutics that have the potential to outrun the virus in the host-virus arms race.

## Author contributions

S.H. and M.M.H.L. conceptualized and designed the study. S.H. and J.G. performed the experiments and analyzed the data. L.N. generated the ePB 3XFLAG ZAPS/L WT single cell clones and provided technical guidance on the RNA binding assay. D.E. curated the alignment of mammalian ZAP, generated the phylogeny tree, and provided expertise in bioinformatics. S.H. wrote the first draft of the manuscript with substantial help from J.G. M.M.H.L. provided critical feedback and S.H. edited subsequent drafts. All authors contributed to manuscript revision, read, and approved the submitted version.

## Acknowledgments

Flow cytometry was performed in the UCLA Jonsson Comprehensive Cancer Center (JCCC) Flow Cytometry Core Facility that is supported by the National Institutes of Health award P30 CA016042 and by the JCCC. RT-qPCR was performed in the UCLA AIDS Institute that is supported by the James B. Pendleton Charitable Trust and the McCarthy Family Foundation.

Molecular structures were performed with UCSF Chimera by the Resource for Biocomputing, Visualization, and Informatics at the University of California, San Francisco (NIH P41-GM103311). This work was funded by NIH grants (R01AI158704; M.M.H.L.), UC Cancer Research Coordinating Committee Faculty Seed Grant (CRN-20-637544; M.M.H.L.), UCLA AIDS Institute and Charity Treks 2019 Seed Grant (M.M.H.L.), and Johanna and Joseph H. Shaper Family Chair (M.M.H.L.).

We thank Dr. Nandita Garud, Dr. Kirk Lohmueller, Dr. Ting-Ting Wu, Erin Kim, Martin Ruvalcaba, and Dr. Zhenlan Yao for their invaluable feedback on the project and critical reading of the manuscript.

## Materials and methods

### Cell culture

HEK293T (parental and ZAP knockout) cells were gifts from Dr. Akinori Takaoka at Hokkaido University (32) and maintained in Dulbecco’s Modified Eagle Medium (DMEM; Thermo Fisher Scientific, Waltham, MA) with 10% fetal bovine serum (FBS; Avantor Seradigm, Radnor, PA). Baby hamster kidney 21 cells (BHK-21; American Type Culture Collection, Manasass, VA) cells were maintained in Minimal Essential Media (Thermo Fisher Scientific) with 7.5% FBS. 0.1mg/mL poly-L-lysine hydrobromide (Millipore Sigma, Darmstadt, Germany) and water were used to coat cell culture dishes when thawing or seeding each cell line to promote cell adhesion and recovery.

### Plasmid

WT or mutant ZAP was cloned into the plasmid pcDNA3.1-3XFLAG (gift from Dr. Oliver Fregoso, University of California, Los Angeles) as previously described (37). 3XFLAG-ZAPS and -ZAPL were amplified from the pcDNA3.1-3XFLAG plasmids using primers to add ClaI and NotI restriction sites for ligation into the ePB vector (gift from Dr. Ali Brivanlou, Rockefeller University) (31). Full-length TRIM25 (gift from Dr. Jae U. Jung at Cleveland Clinic Lerner Research Institute) (49) was cloned into pcDNA3.1-myc as previously described (50). ZAP positive selection mutants were generated by the Q5 Site-Directed Mutagenesis Kit (New England Biolabs, Ipswich, MA) or synthesized as a gene block (Twist Bioscience, South San Francisco, CA) with ClaI and NotI restriction sites and ligated into the ePB vector. The identity of all plasmids was confirmed by Sanger (Genewiz/Azenta, South Plainfield, NJ) and whole-plasmid sequencing (Primordium, Monrovia, CA). See S1 File for a list of all primers used in this study.

### Generation of ZAP inducible cell lines

All ZAP inducible cell lines were made via the ePB transposon system in ZAP KO HEK293T cells. Specifically, ZAP KO HEK293T cells were transfected with equal amounts of the transposase plasmid and an ePB transposon vector containing WT or mutant ZAP using X-tremeGENE9 DNA Transfection Reagent (Roche Life Science, Basel, Switzerland) in Opti-MEM (Thermo Fisher Scientific) following manufacturer’s instructions. 1µg/mL puromycin was added 48 hours post-transfection to select for ZAP KO 293T cells that have incorporated the ePB transposon. Our ZAPS WT and ZAPL WT cell lines were made by selecting single cell clones that follow two criteria: 1) robustly express ZAP following 24 hours of 1µg/mL doxycycline treatment, and 2) recapitulate differential alphaviral sensitivities (S3 Fig) similar to previously generated bulk cell lines with inducible ZAP expression (7,50). The mutant ZAP cell lines in this study were bulk cells that survived after puromycin selection. Comparable inducible ZAP expression in each cell line was validated by immunoblotting following treatment with 1μg/mL doxycycline.

### Viruses and infections

SINV (Toto1101) (51), SINV expressing luciferase (Toto1101/Luc and Toto1101/Luc:ts6) (35), SINV expressing enhanced green fluorescent protein (EGFP) (TE/5’2J/GFP) (52), RRV expressing EGFP (gift from Dr. Mark Heise, University of North Carolina) (53), ONNV expressing EGFP (gift from Dr. Steve Higgs, Kansas State University) (54), CHIKV vaccine strain 181/clone 25 (gift from Scott Weaver, The University of Texas Medical Branch at Galveston) (55) expressing EGFP, and VEEV vaccine strain TC-83 expressing EGFP (gift from Dr. Ilya Frolov, University of Alabama at Birmingham) have been previously described (8,50). All alphaviral stocks were generated and titered in BHK-21 cells (35).

ZAPS/L WT and mutant cell lines were induced for ZAP expression with 1µg/mL of doxycycline 1 day prior to virus infection. To quantify SINV replication, cells were infected with SINV with a luciferase reporter gene (Toto1101/Luc) and harvested 24 hours post-infection. To quantify SINV translation, cells were infected with a replication-deficient temperature-sensitive SINV (Toto1101/Luc:ts6) at 37°C for 1 hour to allow virus adsorption, followed by incubation at 40°C and harvested at the specified timepoints. Harvested lysates were measured for luciferase units following manufacturer’s instructions of the Luciferase Assay System (Promega, Madison, WI). To determine fold inhibition, the average relative luciferase unit (RLU) of +dox was divided by that of -dox for each cell line.

To quantify infection by GFP-alphaviruses, infection was performed as described above and fixed in PBS with 1% FBS and 2% formaldehyde 24 hours post-infection. The fixed cells were analyzed on the Attune NxT Flow Cytometer (Thermo Fisher Scientific), courtesy of the UCLA Flow Cytometry Core. To determine fold change, the GFP percentage of +dox cells in each biological triplicate was divided by the average GFP percentage of -dox cells across biological triplicates for each independent experiment. Then, the fold changes were averaged across the independent experiments.

### Quantification of SINV virion production via plaque assay

To quantify SINV virion production in ZAPL WT or mutant cells, ZAP expression was induced by 1µg/mL doxycycline 1 day prior to infection and infected with SINV Toto1101. The viral supernatant was collected at specific timepoints. To determine viral titers, BHK-21 cells were infected with the viral supernatant at 6 10-fold dilutions and incubated at 37°C for 1 hour with gentle rocking every 15 min. Avicel (RC-581 NF, pharm grade, DuPont Nutrition & Health) overlay consisting of 2X MEM and 4.5% Avicel was added to each well and the plate was incubated at 37°C overnight. On the following day, cells were fixed with 7% formaldehyde for 15 minutes and stained with 1X crystal violet. The plates were washed and the plaques counted after drying.

### Poly(I:C) stimulation, RNA extraction, and quantitative reverse transcription PCR (RT-qPCR)

To stimulate cells with a double-stranded RNA mimic, poly(I:C) diluted in Opti-MEM was incubated with Lipofectamine RNAiMax Transfection Reagent (Thermo Fisher Scientific) before being added to ZAPL WT or mutant cells. 1 day after poly(I:C) stimulation, total RNA was extracted from cells using the Quick-RNA kit (Zymo Research). The amount of RNA template was equalized for reverse transcription using the Protoscript II First Strand cDNA Synthesis Kit (New England Biolabs) and random hexamers. RT-qPCR was performed using 10-fold-diluted cDNA and the Luna Universal qPCR Master Mix (New England Biolabs) in the CFX Real-Time PCR system (Bio-Rad), courtesy of the UCLA Virology Core. qPCR conditions were as previously described (50). Target transcript levels were determined by normalizing the target transcript CT value to the RPS11 transcript CT value. Fold change was calculated using this normalized value relative to that of the corresponding cell line untreated with dox and unstimulated with poly(I:C) (CT method). For RT-qPCR primers, see S1 File.

### Immunoblot analysis

Proteins were visualized using SDS-PAGE with 4-20% Mini-PROTEAN TGX Precast Protein Gels (Bio-Rad) in NuPAGE MOPS SDS Running Buffer (Invitrogen) and transferred to a PVDF membrane (Bio-Rad). The proteins of interest were probed with the corresponding primary and secondary antibodies, followed by visualization on a ChemiDoc imager (Bio-Rad, Hercules, CA) using the ProSignal Pico ECL Reagent detection reagent (Genesee Scientific, El Cajon, CA).

Primary antibody 1:20,000 anti-FLAG (Sigma-Aldrich), 1:20,000 anti-actin-HRP (Sigma-Aldrich), or 1:1000 anti-poly(ADP-ribose) (Abcam), and secondary antibody 1:20,000 goat anti-mouse HRP (Jackson ImmunoResearch, West Grove, PA), or 1:20,000 goat anti-rabbit HRP (Thermo Fisher Scientific) were used to probe the protein of interest. See Table S2 for more detail.

### *In vitro* biotinylation of SINV RNA and RNA pulldown assay

The genomic SINV DNA template was digested by XhoI and *in vitro* transcribed using SP6 RNA polymerase (New England Biolabs) and 0.5mM biotin-16-UTP (Roche Life Science, Penzberg, Germany) as previously described (37). RNA biotinylation was confirmed by streptavidin-HRP dot blot as previously described (8).

*In vitro* RNA pulldown was performed as previously described (37). ZAP expression was induced in ePB ZAP cell lines and the protein lysates were harvested in CHAPS buffer (10mM Tris-HCl pH7.5, 1mM MgCl_2_, 1mM EDTA, 0.5% CHAPS, 10% glycerol, 5mM beta-mercaptoethanol, and protease inhibitor) 24 hours later. 0.4pmol of biotinylated SINV RNA was incubated with normalized amounts of protein lysates and RNA binding buffer containing RNaseOUT (Thermo Fisher), heparin (Sigma-Aldrich), and yeast tRNA (Thermo Fisher) to minimize non-specific binding. The lysate-RNA samples were incubated with Dynabeads M-280 Streptavidin (Invitrogen) on a shaker for 30 min at room temperature. Protein visualization on a ChemiDoc imager was as described above.

### Immunoprecipitation assays

To test interaction with TRIM25, ZAP KO HEK293T cells were transfected with pcDNA3.1-3XFLAG-ZAPL and pcDNA3.1-myc-TRIM25. Cells were lysed in FLAG buffer (100mM Tris HCl pH8.0, 150mM NaCl, 5mM EDTA, 5% glycerol, 0.1% NP-40, 1mM DTT, and protease inhibitor) and incubated on a rotator at 4°C for 30 min. After equilibration, FLAG beads were incubated with lysates on a rotator at 4°C for 45 min. Immunoprecipitated samples were washed 3 times with FLAG buffer and eluted in Laemmli buffer for immunoblotting.

PAR binding assay was based on (17) with modification. Briefly, ZAP inducible cells were lysed in lysis buffer containing 50mM Tris-HCl pH7.5, 150mM NaCl, 0.2% Triton X-100, protease inhibitor, and 1µM PARG inhibitor PDD 00017273 (Tocris Bioscience, Bristol, UK). After equilibration, FLAG beads were incubated with lysates on a rotator at 4°C for 1 hour and 30 min. Bound lysates were washed 3 times with IP buffer (50mM Tris-HCl pH7.5, 150mM NaCl, and 0.2% Triton X-100) and eluted in Laemmli buffer for immunoblotting.

### Sequence alignment, phylogenetic tree, and positive selection analysis

The coding sequence (CDS) of human ZAPXL was used to search for orthologs in 260 other mammalian genome assemblies with a contig size of at least 30kb in the NCBI assembly database as of July 2020 to minimize truncated orthologous coding sequences. To extract the orthologous coding sequences of ZAP, we used best Blat reciprocal hits from the human CDS to every other mammalian genome, and back to the human genome (matching all possible reading frames, minimum identity of 30%, and the “fine” option activated).

The 261 orthologous ZAP were aligned to human ZAPXL with MACSE v2 with maximum accuracy settings (S2 File). The alignments generated by MACSE v2 were then cleaned by HMMcleaner using default parameters to remove errors from genome sequencing and “false exons” that might have been introduced during the Blat search. Visual inspection confirmed that the resulting alignment had a very low number of visibly ambiguous or erroneous segments.

The phylogenetic tree of the 261 mammals was built using IQ-Tree to generate the consensus, maximum likelihood tree with a GTR substitution model with six parameters (GTR-6) which provided the best fit (S2 File).

More complete details on the alignment and phylogenetic tree reconstruction are given in (56) as the same exact pipeline was used for this study.

The positive selection analyses FEL, MEME, and FUBAR were performed using HyPhy from the command line, with the aforementioned alignment and mammalian tree as inputs. Rodrigue *et al.*’s positive selection test based on a Mutation-Selection balance (Mutselomega) was used as described in (27). Briefly, Mutation-Selection balance tests attempt to provide higher statistical power to detect positive selection by better accounting for selective constraint in coding sequences, beyond the usual arbitrary use of the dN/dS>1 threshold by other selection tests.

### Statistical analysis

Experiments were performed at least two independent times and statistical analyses were performed on biological replicates from triplicate wells using GraphPad Prism.

## Supporting information

**S1 Fig. Positive selection and domains of ZAP.**

(A) Positive selection analyses on ZAPXL of 261 mammalian species detected by the FEL, MEME, FUBAR, and Rodrigue methods. (B) ZAP isoforms annotated with their domains. The four ZAP splice variants are depicted here: ZAPS (short), ZAPM (medium), ZAPL (long), and ZAPXL (extra-long). All isoforms contain the zinc finger (Z1-Z5, pink) and WWE domains (green), but only ZAPXL and ZAPL have a catalytically inactive PARP-like domain (indigo). ZAPXL and ZAPM also share an extended exon 4 (teal). The amino acid numbering of domains is based on (6) and (7).

**S2 Fig. N658A mutant induces interferon (IFN) and interferon-stimulated gene (ISG) levels similar to WT.**

ZAPL WT or N658A inducible ZAP KO 293T cells were untreated, treated with poly(I:C), or treated with both poly(I:C) and dox. RNA was harvested for RT-qPCR. mRNA levels of IFN or the ISG IFIT1 in each condition were normalized to that of the respective cell line without poly(I:C) and without dox. Data are combined from two independent experiments. Asterisks indicate statistically significant differences as compared to the -dox -poly(I:C) condition for each cell line (Two-way ANOVA and Tukey’s multiple comparisons test: *, p<0.05; **, p<0.01; ****, p<0.0001).

**S3 Fig. Characterization of WT ZAP inducible single clone cell lines.**

(A) Western blot of ZAPS and ZAPL WT inducible ZAP KO 293T cell lysates. Each single clone cell line was treated with dilutions of doxycycline 24 hours after seeding. Cell lysates were harvested 24 hours after dox treatment. (B) ZAPS and ZAPL WT inducible ZAP KO 293T cells were induced for ZAP expression 24 hours before infection by GFP-expressing alphaviruses and harvested at the time listed for flow cytometry (SINV, MOI = 10, harvest 8 h.p.i.; RRV, MOI = 10, harvest 24 h.p.i.; ONNV, MOI = 0.1, harvest 18 h.p.i.).

**S1 File. Primers used in this study.**

**S2 File. Alignment and phylogenetic tree from the 261 mammalian ZAP sequences.**

